# CD8⁺ T cells induce interstrand crosslinking-associated DNA damage in neurons

**DOI:** 10.64898/2026.03.14.711737

**Authors:** Britanie M. Blackhurst, Aayush Bhatt, Emily Kretchmer, Autumn E. Tucker, Bree Kurtz, Katie L. Reagin, Kristen E. Funk

**Affiliations:** Department of Biological Sciences, University of North Carolina at Charlotte, Charlotte, North Carolina, USA; Burnett School of Biomedical Sciences, University of Central Florida College of Medicine, Orlando, Florida, USA

**Keywords:** DNA damage, Fanconi anemia repair pathway, Interstrand crosslinking, Kunjin virus, Neuroinflammation, Neurons, T cells, Viral encephalitis

## Abstract

Viral pathogens cause neurologic sequelae during acute and post-acute phases of infection. CD8^+^ T cells are hypothesized to contribute to these effects, but the mechanisms through which they act are poorly understood. We posited that viral infections and/or antiviral immune responses induce DNA damage, which may underlie neuronal dysfunction. Using a model of neurotropic flavivirus infection, we found that genes associated with interstrand crosslinking (ICL) DNA damage were upregulated post-infection, temporally congruent with T cell infiltration. Using an *in vitro* co-culture system, our results demonstrate that CD8^+^ T cells induced ICL-like damage in primary neurons, independent of antigen-specific interactions or direct contact. Human transcriptomic data also showed overexpression of genes associated with ICL damage in the brains of people with Parkinson’s disease, Alzheimer’s disease, and multiple sclerosis, which are neurologic diseases characterized by neuroinflammation. Together, these data indicate that CD8^+^ T cells cause genotoxic DNA damage in neurons, which may underlie the neurologic dysfunction seen in neurodegenerative conditions.

**Summary:** Results indicate that CD8^+^ T cells induce interstrand crosslinking-like DNA damage in neurons independent of antigen-specificity in a mouse model of viral infection, *in vitro* primary cell culture system, and human neurologic diseases. These findings provide insight on the mechanistic connection between neuroinflammation and neurologic dysfunction.

## Introduction

Viral infections are associated with neurologic sequelae that include cognitive decline, motor dysfunction, microcephaly, ocular manifestations, and Guillan-Barré syndrome (Blackhurst and Funk, 2023). Neurotropic pathogens can cause neuronal dysfunction by inducing cell death, mitochondrial damage, protein misfolding and demyelination, among other mechanisms (Blackhurst and Funk, 2023); however, peripheral infections, in which virus does not infect neurons or the central nervous system (CNS) directly, can also cause neurologic sequelae due to metabolic dysfunction, blood vessel damage, and immunopathology (Valerio et al., 2021). Post-infectious neurologic dysfunction has been attributed to the immune system for a number of infectious diseases including West Nile virus (WNV) (Garcia et al., 2014), Influenza A virus (Nath and Kolson, 2025), and SARS-CoV-2 (Talla et al., 2023), suggesting an important role of the immune system in contributing to post-infectious cognitive dysfunction.

Inflammatory events, such as infections, stroke, neurodegeneration, injury, and even aging, cause sentinel cells such as microglia and astrocytes to become activated and secrete cytokines and chemokines that open the blood-brain barrier and recruit peripheral immune cells to the CNS, including T cells, which are normally excluded during homeostatic conditions (Yang et al., 2022; Li et al., 2025). The recruitment of CD8^+^ T cells, known as “cytotoxic T cells,” during neurotropic viral infection is critical for the clearance of viral reservoirs from the brain and host survival (Shrestha and Diamond, 2004; Shrestha et al., 2006, 2012). Maximal CD8⁺ T cell activation and clonal expansion typically requires three signals: (1) TCR recognition of peptide–MHC I, (2) co-stimulation via CD28 engagement by CD80/CD86 on antigen-presenting cells, and (3) inflammatory cytokines, most prominently Interleukin (IL)-12 and type I interferons (IFN-α/β) that promote proliferation, survival, and effector differentiation (Keppler et al., 2012). However, CD8^+^ T cells can be partially activated independent of antigen-TCR engagement by cytokine signaling alone, known as “bystander activation” (Lamichhane et al., 2019; Watson et al., 2024). Cytokines including IL-12, IL-18, and Tumor Necrosis Factor (TNF) can promote this bystander activation, which provides innate immune protection against viruses and bacteria by promoting expression of CD69 and IFN-(Freeman et al., 2012; Raué et al., 2004; Lertmemongkolchai et al., 2001).

T cells are normally excluded from the brain during homeostatic conditions, but CD8^+^ T cells are recruited to the brain by coordinated chemokine signaling from neurons and resident immune cells, including microglia and astrocytes. In response to viral infection, neurons secrete CXCL10, which is the binding partner of CXCR3 expressed by activated CD8^+^ T cells (Klein et al., 2005; Campanella et al., 2008).

Additional signaling across the IL-1-CXCL12-CXCR4 axis promotes the migration of T cells across the blood-brain barrier into the brain parenchyma (McCandless et al., 2008; Durrant et al., 2014). However, recent data show that neurotropic infection is not specifically required to drive CD8^+^ T cells into the brain (Steinbach et al., 2016a). In fact, microbial exposure is sufficient to promote brain-penetrant T cell infiltration, and the resulting brain-resident memory T cells are often specific for peripheral antigens rather than CNS pathogens (Mix et al., 2025). In support of this, our prior research showed an abundance of CD8^+^ T cells recruited to the brain of aged mice during infection that were not specific to the immunodominant viral antigen (Reagin et al., 2024).

There is a growing body of evidence indicating that CD8^+^ T cells negatively impact neuronal function and brain health (Reagin and Funk, 2022). We have previously shown that antigen non-specific CD8^+^ T cells can induce neuronal death *in vitro*, but the molecular and cellular mechanisms through which this occurred was not examined (Reagin et al., 2024). When CD8^+^ T cells encounter an MHC-I complex loaded with their cognate peptide antigen, they can induce neuronal apoptosis through the release of perforin and TRAIL, as well as Fas/Fas ligand interactions (Shrestha et al., 2006, 2012; Shrestha and Diamond, 2007). In addition to direct effects, CD8^+^ T cells cause indirect effects through the release of proinflammatory cytokines, notably IFN-γ, TNF, and IL-6, that orchestrate responses in both resident and infiltrating immune cells (Tsunoda et al., 2006; Kaya et al., 2022). .

Activated CD8^+^ T cells also secrete reactive metabolites that may compromise neuronal genomic stability. For example, during heightened glycolytic metabolism, activated CD8⁺ T cells release reactive oxidative species (ROS) and reactive nitrogen species (RNS), which can cause oxidative DNA lesions such as 8-oxoguanine, abasic sites, and double-strand breaks (Szabó and Ohshima, 1997; Sena et al., 2013). Even antigen-experienced CD8⁺ T cells that are not fully reactivated, often marked by low CD44 expression and reduced glycolytic flux, can exert neuropathology via activation of the Natural Killer cell receptor (Balint et al., 2024) and fatty-acid oxidation (O’Sullivan et al., 2014). These bystander CD8⁺ T cells generate lower levels of ROS and RNS but display increased lipid peroxidation, leading to the production of reactive aldehydes such as 4-hydroxynonenal (4-HNE) and malondialdehyde (MDA), shown to cause interstrand crosslink (ICL) DNA damage formation (Xu et al., 2021). These aldehydes can diffuse into the parenchyma and induce ICL DNA damage in nearby neurons with deficient DNA repair mechanisms due to their post-mitotic nature (Carnie et al., 2024). The effect of persistent CD8⁺ T cell activity in the brain on immune regulation, synapses, and behavior has now been well described (Zinselmeyer et al., 2013; Steinbach et al., 2016b; Galiano-Landeira et al., 2020), but its impact on neuronal DNA integrity and its contribution to cumulative neurodegeneration remains largely unexplored.

This study aimed to determine the impact of bystander CD8^+^ T cells on neurons, specifically on DNA damage. Our results indicate that brain-infiltrating CD8^+^ T cells induce highly genotoxic ICL-like DNA damage in neurons. We show this in a mouse model of neurotropic flavivirus infection and using an *in vitro* co-culture system of primary mouse neurons and CD8^+^ T cells. Using a transwell system we show that DNA damage is mediated by a soluble factor produced by both stimulated and unstimulated CD8^+^ T cells rather than direct antigen-specific interactions. We then analyzed publicly available human transcriptome datasets from individuals with Parkinson’s disease (PD), Alzheimer’s disease (AD), and multiple sclerosis (MS), and our findings support a role for ICL-associated DNA damage/repair pathways in these human neurologic diseases. These results provide new evidence that CD8^+^ T cells impact neurons through ICL-like DNA damage and, specifically, a mechanism by which bystander activated CD8^+^ T cells affect uninfected neurons, potentially contributing to neurodegenerative diseases.

## Results

### Kunjin virus infection upregulates ICL-associated DNA damage pathways in a mouse model of viral encephalitis

To establish whether neurotropic viral infection causes DNA damage, we used a mouse model of neurotropic viral infection. We used a naturally attenuated strain of WNV, Kunjin virus (KUNV), to infect wildtype C57BL/6J mice intracranially (i.c.). To confirm that this inoculation paradigm resulted in productive infection in the CNS, we measured viral titers in the hippocampus (HC) and cortex (CX) (Fig. 1 A, B), as well as the olfactory bulb (OB), cerebellum (CB), and brainstem (BS) (Fig. S1 A-C) at specific days post-infection (d.p.i.). Results show that replicating virus was detected in the brain starting at 6 d.p.i., peaked at 6–8 d.p.i., and declined substantially by 12 d.p.i., consistent with what has been reported previously using the laboratory-attenuated WNV-NS5-E218A strain (Vasek et al., 2016). As expected, approximately 40% of mice succumbed to infection (Fig. S1 D). Mice that survived infection began regaining lost weight around 20 d.p.i. and recovered to their initial weight around 28 d.p.i. (Fig. S1 E).

**Figure 1.**
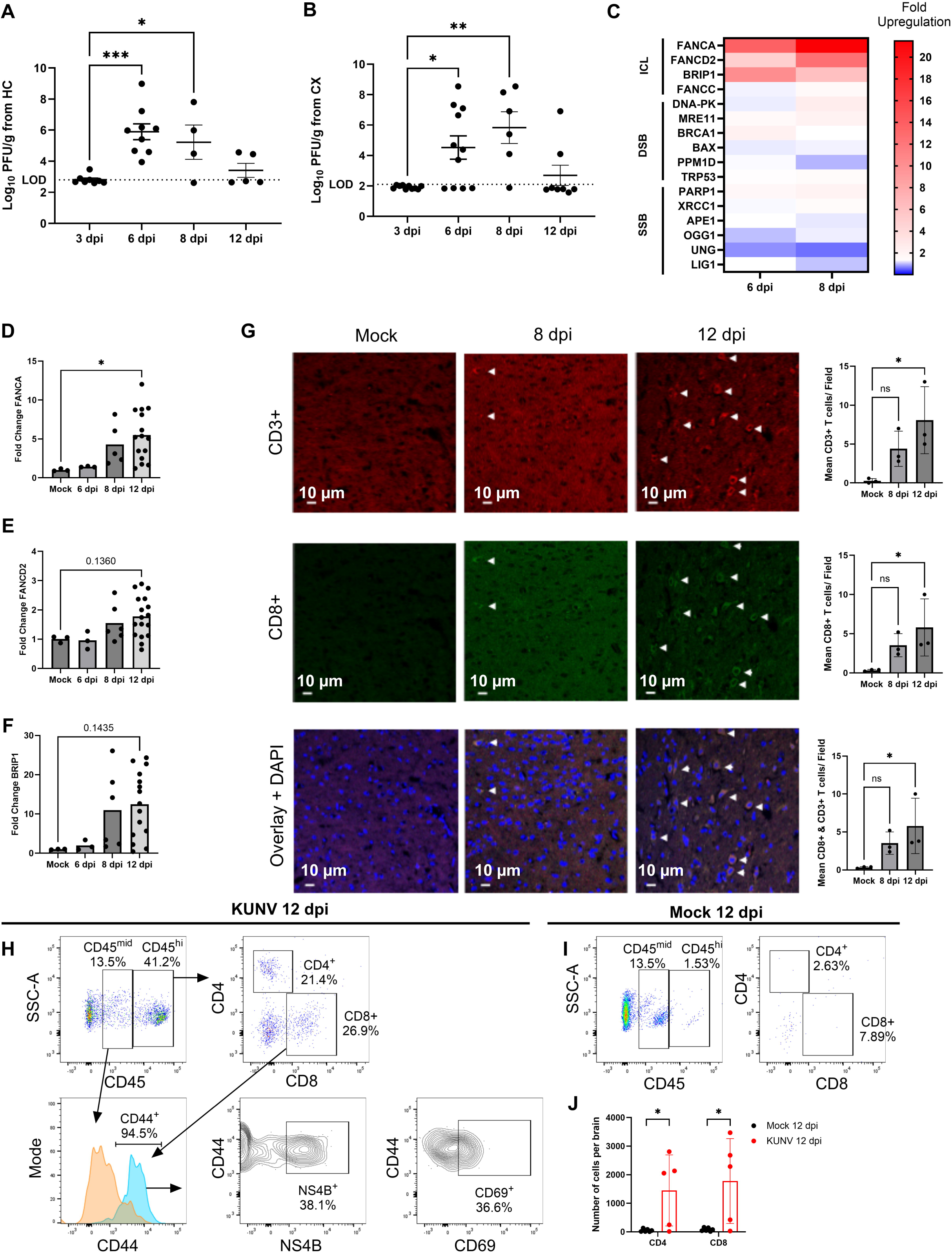
ICL gene expression is upregulated in mouse model of viral encephalitis, temporally congruent with CD8^+^ T cell infiltration. Viral titers of (A) hippocampal (HC) and (B) cortical (CX) tissue at 3, 6, 8 and 12 d.p.i. (C) Heatmap of gene expression from RT-qPCR of genes associated with ICL, DSB and SSB DNA damage depicted by fold change of n = 2 murine hippocampus at 6 d.p.i. and 8 d.p.i. Gene expression changes in ICL DNA damage genes (D) *FANCA*, (E) *FANCD2*, and (F) *BRIP1* measured by RT-qPCR from mock infected, 6, 8 and 12 d.p.i. murine hippocampal tissue. (G) Representative IHC images of cortical brain region from mock, 8 d.p.i., and 12 d.p.i. with arrows pointing to CD8^+^ T cells (CD3^+^, CD8^+^ and DAPI). Images were captured at 20x magnification. Quantification of number of CD3^+^, CD8^+^ and double-positive cells per imaging field are shown to right. Representative flow cytometry plots of cortical tissue collected from (H) KUNV-infected (n=3M/3F) and (I) mock-inoculated (n=3M/3F) mice at 12 dpi. Virally infected plots show workflow to identify CD4^+^ and CD8^+^ T lymphocytes, CD44^+^ antigen-educated cells, NS4B^+^ KUNV-specific CD8^+^ T cells and CD69^+^ CD8^+^ T cells, which are absent from mock-inoculated brains. (J) Quantification of the number of CD4^+^ and CD8^+^ T cells in cortices of mock-and KUNV-inoculated mice. Statistical analysis by 1way ANOVA with multiple comparisons. *, P < 0.05; **, P < 0.01; ***, P <0.001.

To determine whether viral infection impacted genome integrity, we quantified expression of DNA damage response genes across major repair pathways. Genes specifically associated with ICL repair, including *FANCA, FANCD2,* and *BRIP1*, were selectively upregulated in the HC at 6-8 d.p.i. in a small number of mice (Fig. 1 C). In contrast, genes primarily connected to double-strand break (DSB) or single-strand break (SSB) repair exhibited little or no induction, indicating a preferential activation of the ICL repair pathway rather than a generalized DNA damage response following KUNV infection. We confirmed significant upregulation of *FANCA*, a core component of the fanconi anemia response complex, and a trending upregulation of *FANCD2* and *BRIP1*, also involved in ICL repair, in a larger cohort of mice at 12 d.p.i. in the HC (Fig. 1 D-F). We noted that the most significant upregulation of *FANCA, FANCD2*, and *BRIP1* occurred after viral titers had largely cleared from the HC, temporally congruent with expected T cell infiltration into the infected CNS, suggesting that ICL-associated signaling may be due to antiviral immune responses rather than direct effects of the virus.

To assess T cell infiltration in this model, we performed immunohistochemical (IHC) analysis for T cell markers CD3 and CD8 on frozen brain sections from KUNV-infected mice at 8 and 12 d.p.i.. Analysis revealed a significant increase in CD3⁺ and CD8⁺ T cells in the cortical parenchyma at 12 d.p.i. versus mock controls (Fig. 1 G). To further assess T cell infiltration and activation status, we performed flow cytometry analysis on leukocytes isolated from the cortices of mock or KUNV-infected mice at 12 d.p.i. (Fig. 1 H-J). Significantly more CD4^+^ and CD8^+^ T cells were present in the brains of mice post-KUNV-infection compared with mock-infected controls (Fig. 1 J). As is typical for brain-infiltrating CD8^+^ T cells, more than 90% of cells express CD44, a cell surface glycoprotein that is upregulated upon TCR engagement and involved in T cell homing and extravasation (DeGrendele et al., 1997). Of the CD44^+^CD8^+^ T cells, 35-40% were specific to the immunodominant KUNV NS4B tetramer peptide or express early activation marker CD69^+^, indicating that both KUNV-specific and bystander CD8^+^ T cells enter the brain at this timepoint (Fig. 1 H). We hypothesized that both antigen-specific and bystander CD8^+^ T cells can induce ICL DNA damage in neurons, and sought to test whether this damage may be incurred via direct interactions, indirect interactions, or both.

### Direct co-culture with CD8⁺ T cells induce neuronal ICL-type DNA damage

To test whether CD8⁺ T cells are sufficient to induce ICL-associated DNA damage responses in neurons, we established an *in vitro* co-culture system using murine primary neurons and splenic CD8⁺ T cells. This approach was motivated by prior work showing that CD8⁺ T cells can trigger neuronal injury *in vitro* even in the absence of cognate antigen recognition (Reagin et al., 2024), allowing for isolation of T cell-derived effector mechanisms apart from virus- or antigen-specific variables. Splenic CD8⁺ T cells were bulk-stimulated using PMA/ionomycin or were left unstimulated, to model activated versus naïve states. Compared with neurons cultured in isolation, neurons exposed to either unstimulated (naive) or PMA/ionomycin-stimulated (activated) CD8⁺ T cells exhibited significant upregulation of *FANCA*, *FANCD2*, *BRIP1*, and *FANCC* (Fig. 2A–D), recapitulating the transcriptional signature observed in mice post-infection.

**Figure 2.**
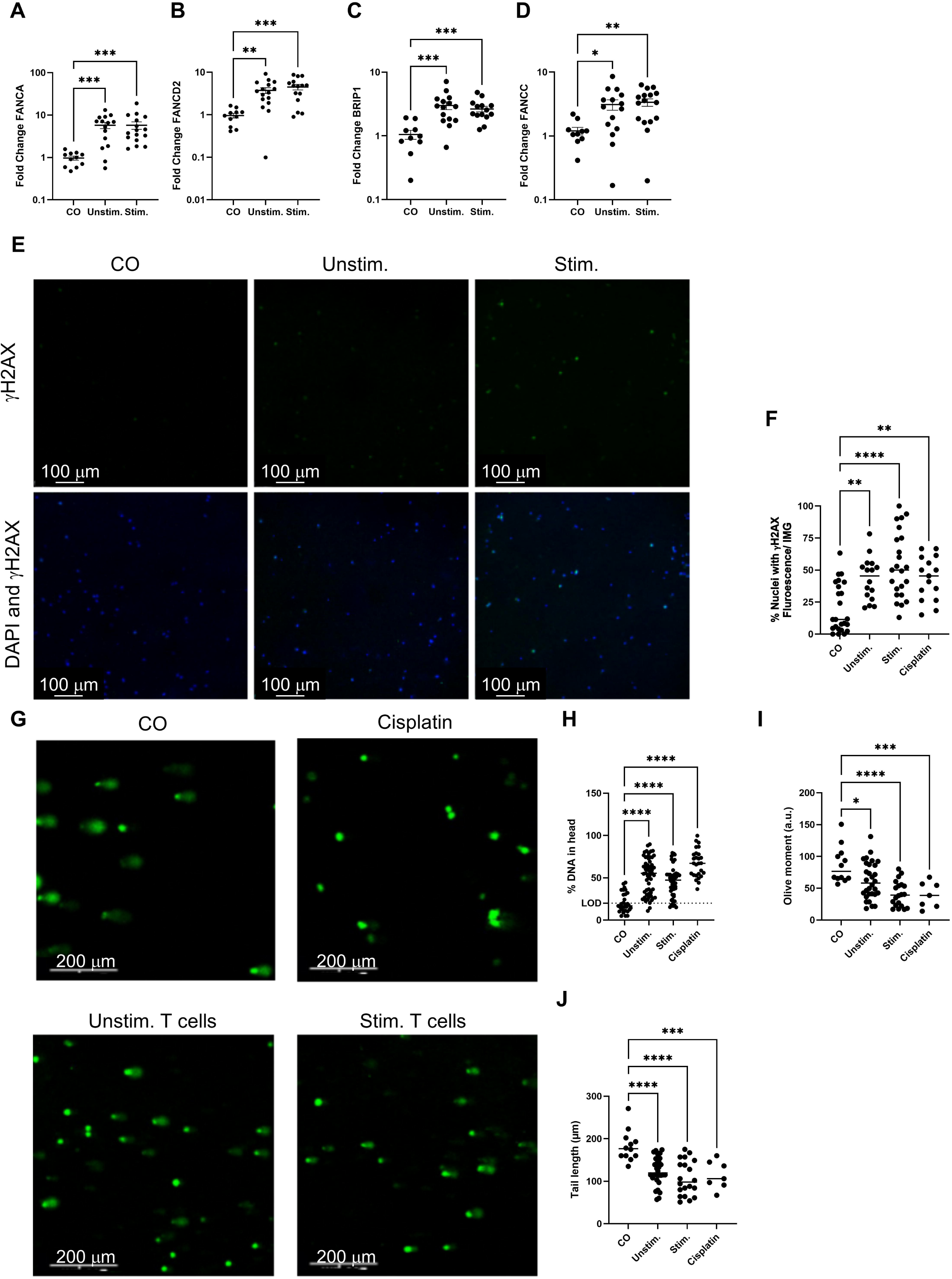
Primary CD8^+^ T cells co-cultured with primary neurons induce ICL DNA damage. Primary neurons were cultured alone (CO), or co-cultured with splenic CD8^+^ T cells that were unstimulated (Unstim.) or stimulated (Stim.) with PMA/ionomycin. Neuronal gene transcription was measured by RT-qPCR of (A) *FANCA*, (B) *FANCD2,* (C) *BRIP1*, and (D) *FANCC*. (E) Representative ICC images of primary neurons stained for γH2AX (green) and counterstained with DAPI (blue). (F) Total number of γH2AX positive nuclei divided by the number of total nuclei per ICC image in control embryonic neurons in the treatment groups listed in 2A or treated with Cisplatin, a positive control for inducing ICL-damage. (G) Representative images of comets from nuclear DNA from neurons cultured alone, with ICL inducer cisplatin, or co-cultured with unstimulated/stimulated CD8^+^ T cells. (H-J) Quantification of (H) percent DNA in comet head, (I) olive moment, and (J) tail length of comets, counted per single nucleus in cultured neurons. Statistical analysis by Kruskal-Wallis (A-D) and ordinary 1way ANOVA with multiple comparisons (F, H-J) . *, P < 0.05; **, P < 0.01; ***, P <0.001.

As gene upregulation does not directly indicate DNA damage, we used two more direct measures of breakage: γH2AX foci and comet assay. Phosphorylation of the histone protein H2AX at Serine-139 is specifically termed γH2AX and denotes double-stranded DNA breaks with individual breakage sites distinguishable by ICC (Ivashkevich et al., 2011). ICC quantification revealed increased γH2AX foci in neurons exposed to either unstimulated or stimulated CD8⁺ T cells (Fig. 2 E and F), further supporting the hypothesis that CD8+ T cells induce DNA damage. However, γH2AX does not discriminate between ICLs and other types of DNA lesions such as DSBs and SSBs. To specifically probe for ICLs, we performed a modified alkaline comet assay in which nuclei were treated with H_2_O_2_ to induce DNA breaks, leaving only covalent bonds in the DNA backbone and crosslinked DNA intact. When subjected to electrophoresis, large, crosslinked DNA is visualized as a “comet head” while the smaller, broken DNA is visualized as the “comet tail” (Wu and Jones, 2012). Neurons co-cultured with CD8^+^ T cells demonstrated crosslinked DNA, as evidenced by a greater proportion of DNA in the comet head, reduced tail length, and increased olive moment relative to control neurons cultured alone but subjected to the same H_2_O_2_ and alkaline treatment conditions (Fig. 2 G–J). These results were akin to neurons treated with positive control agent cisplatin, a cancer chemotherapeutic known to induce intrastrand DNA adducts primarily in the form of interstrand crosslinks (Takahara et al., 1995). To test for DNA breaks rather than crosslinks, we performed a standard comet assay in which H_2_O_2_ and alkaline treatment conditions were omitted. These results likewise showed significant differences between the proportion of DNA in the comet head and olive moment when neurons were cultured alone versus co-cultured with either unstimulated or stimulated CD8^+^ T cells, indicating DNA breaks in co-cultured neurons. However, the overall DNA migration was reduced due to the absence of H_2_O_2_ treatment, resulting in lower separation between the two groups and a less sensitive readout (Fig. S2 A–D).

Together, these data show that CD8^+^ T cells induce bona fide DNA crosslinks and to a lesser extent, double and/or single-stranded breaks, in co-cultured neurons.

To confirm that the apparent ICL damage was not simply an artifact of the culture conditions, we co cultured neurons with TK1 cells, an immortalized CD8⁺ T cell line.

While TK1 cells are of CD8^+^ T lymphoblast lineage, they are highly proliferative and express high levels of lymphocyte integrins important for lymphocyte homing (Rüegg et al., 1992; Hu et al., 1992). Thus, TK1 cells are generally useful for studying CD8^+^ T cell migration rather than cytotoxic functions typical of a terminally differentiated CD8^+^ T cell (Uhlemann et al., 1997). Results showed that unlike primary CD8⁺ T cells, TK1 cells failed to induce *FANC* gene upregulation (Fig. S2 E and F), supporting our hypothesis that ICL DNA damage is due to direct or indirect cytotoxic effects of CD8^+^ T cells.

### CD8⁺ T cells induce ICL-associated DNA damage through a contact-independent mechanism

To further examine the mechanism by which this DNA damage occurs and to test whether the effect depends on direct contact versus indirect soluble mediators, we used a transwell system with a 1 µm pore size, which allows co-cultured neurons and CD8^+^ T cells to share growth media and secreted factors but prohibits CD8^+^ T cell transmigration and direct contact (Wolf et al., 2013). Similar to direct contact conditions, neurons exposed to CD8⁺ T cell-conditioned media via transwell inserts exhibited robust induction of *FANCA*, *BRIP1*, and *FANCC* gene expression (Fig. 3 A–D). However, *FANCD2* was not significantly upregulated at the transcript level under transwell conditions. We again sought to confirm direct evidence of DNA damage by γH2AX staining and comet assay. Neurons exposed to transwell CD8⁺ T cells displayed increased γH2AX staining (Fig. 3 E and F) and a comet assay profile indicative of ICLs with elevated percent DNA in the head, reduced tail length, and increased olive moment (Fig. 3 G–J). This evidence of DNA damage was seen from cultures with both PMA/ionomycin stimulated CD8+ T cells and unstimulated CD8+ T cells, suggesting that unstimulated T cells were sufficient to induce damaging effects via indirect contact. In an effort to identify the soluble factor responsible for mediating these effects, we performed a multiplex array to measure the concentration of specific cytokines in the conditioned media from neurons alone or co-cultured with unstimulated/stimulated primary CD8^+^ T cells (Fig. S3 A) or with unstimulated/stimulated TK1 cells (Fig. S3 B). Results show that PMA/ionomycin stimulation increased expression of TNF, IFNγ, IL-2, IL-4, IL-6, IL-13, and IL-17, as expected, in primary CD8^+^ T cells, but not the TK1 cell line; however, we saw no detectable difference in expression of any cytokines tested in media from unstimulated CD8+ T cells compared with neuron alone-conditioned media (Fig. S3). These results confirm that our “unstimulated” culture conditions did not cause T cell activation and eliminate several likely candidates, including TNF, IFNγ, and IL-1β, as causative agents; however, the identity of the agent mediating DNA damage in neurons remains elusive. Altogether, these data demonstrate that physical interaction between CD8^+^ T cells and neurons is not required for ICL induction and instead indicate that CD8⁺ T cells secrete a soluble factor capable of generating ICL-associated DNA damage in bystander neurons.

**Figure 3.**
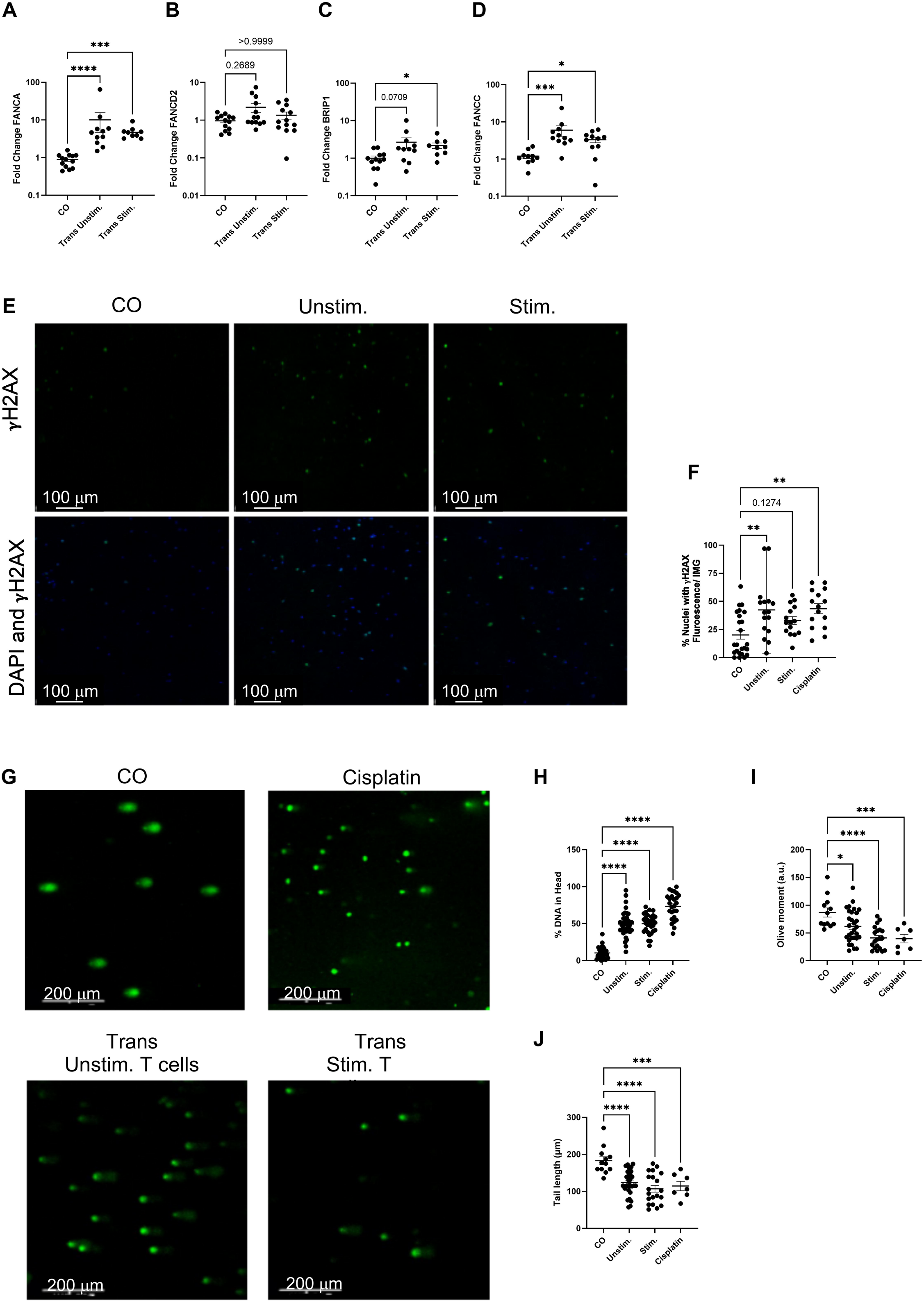
Direct interactions are not necessary for CD8^+^ T cells to induce ICL DNA damage in co-cultured neurons. Primary neurons were cultured alone (CO) or co-cultured using a transwell system with unstimulated (Trans Unstim.) or PMA/ionomycin-stimulated (Trans Stim.) splenic CD8^+^ T cells. Neuronal gene transcription was measured by RT-qPCR of (A) *FANCA*, (B) *FANCD2,* (C) *BRIP1*, and (D) *FANCC*. Results are shown as fold change relative to CO using the 2^-ΔΔCt^ method. (E) Representative ICC images of primary neurons stained for γH2AX (green) and counterstained with DAPI (blue). (F) Total number of γH2AX positive nuclei divided by the number of total nuclei per ICC image for neurons cultured alone, with unstimulated/stimulated T cells, or treated with ICL-inducing positive control cisplatin. (G) Representative image of comet assay on nuclear DNA from primary neurons. (H-J) Quantification of (H) percent DNA in comet head, (I) olive moment, and (J) tail length of comets in primary neurons treated as indicated. Statistical analysis by Kruskal-Wallis (A-D) and ordinary 1way ANOVA with multiple comparisons (F, H-J) . *, P < 0.05; **, P < 0.01; ***, P <0.001.

### ICL-associated DNA damage signatures are elevated in human neurologic disorders

To evaluate whether the ICL-like damage response gene signature that we found in our mouse model of infection extends to human neurologic diseases, we analyzed transcriptomic datasets from patients with PD, AD, and MS, described in Table 1 (Dunckley et al., 2006; Han et al., 2012; Moran et al., 2006). These conditions exhibit diverse neuropathologies, but they are all associated with neuroinflammation and CD8^+^ T cell infiltration (Martirosyan et al., 2024; Zhu et al., 2024; Sulzer et al., 2017; van Nierop et al., 2017; Gate et al., 2020; Lueg et al., 2015; Babbe et al., 2000). Across all three disorders, *FANCA* was significantly upregulated compared with matched controls (Fig. 4 A–C). Supplemental analysis also demonstrated increased Fanconi pathway gene induction, including *FANCD2*, *BRIP1*, and *FANCC* to varying degrees across datasets and brain regions, but particularly in the PD cohort (Fig. S4 A–E), indicating broad activation of ICL repair pathways in human neurologic diseases. The reproducibility of this pathway-level induction across independent cohorts, platforms, brain regions, and disease contexts suggests that Fanconi/ICL-repair activation is a recurring feature of human neuroinflammatory disease.

**Figure 4.**
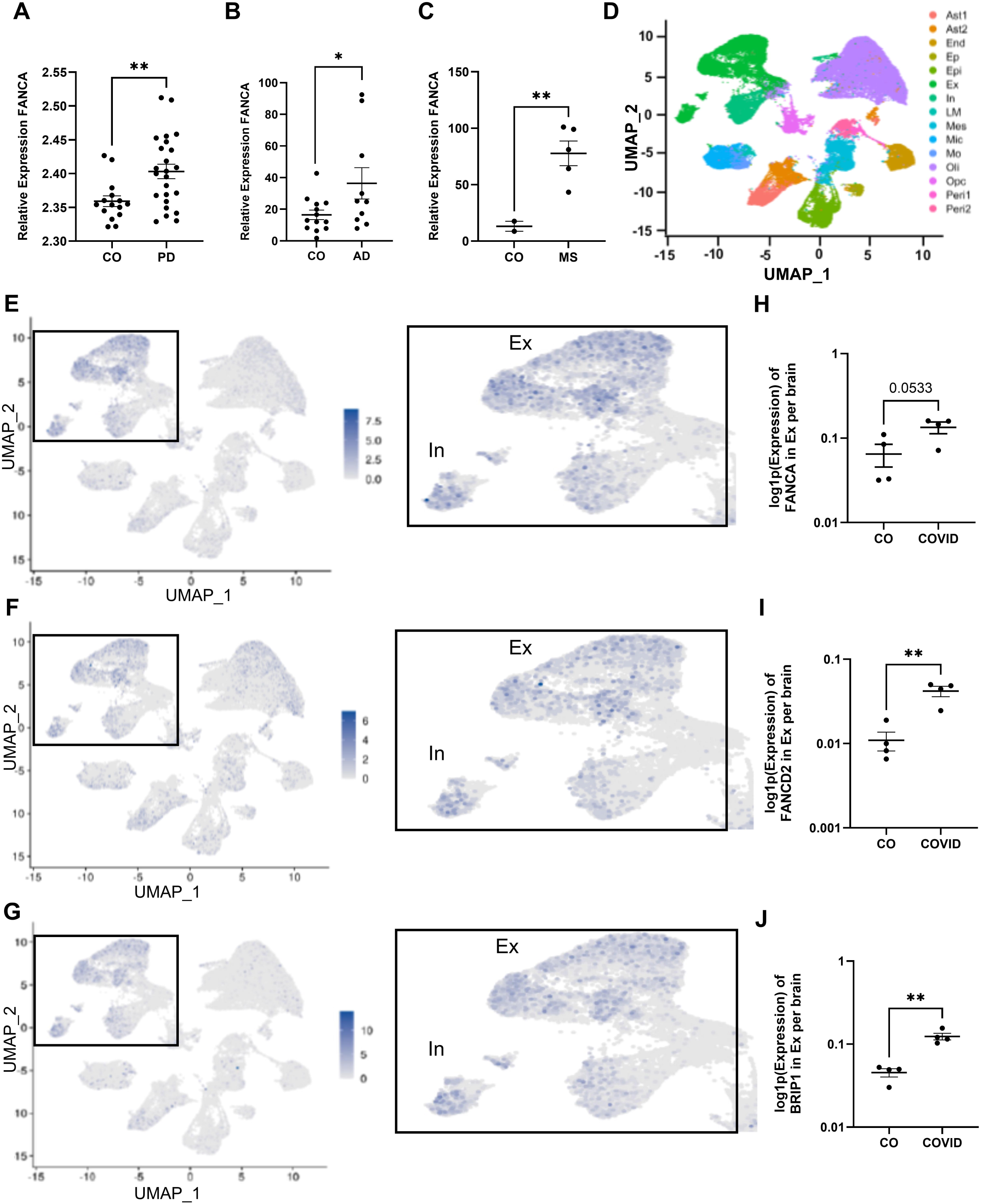
Fanconi anemia pathway transcript expression in human neurologic conditions. Transcriptional RNA expression was examined in human datasets described in Table 1. *FANCA* RNA expression in (A) the medial substantia nigra of PD patients vs cognitively normal controls, (B) microdissected neurons isolated from Alzheimer’s disease brains, with or without neurofibrillary tau pathology, and (C) white matter from MS lesions vs healthy white matter control. (D) UMAP projections distinguishing cell types from snRNA-seq data from the brains of patients who died of severe COVID-19 and non-infected controls. (E-G) UMAP projections of snRNA-seq data from COVID-19 patients highlighting cellular expression of (E) *FANCA*, (F) *FANCD2* and (G) *BRIP1* in excitatory (Ex) and inhibitory (In) neurons. (H-J) Quantification of gene expression in excitatory neurons for (H) *FANCA,* (I) *FANCD2*, and (J) *BRIP1*. Statistical analysis by Welch’s T-test. *, P < 0.05; **, P < 0.01; ***, P <0.001.

**Table 1.** Human disease datasets analyzed.

| Disease of interest | Geo Accession Number | Experimental design | Original publication |
| --- | --- | --- | --- |
| Parkinson's disease | GSE8397 | Microarray analysis of post mortem brain tissue samples from substantia nigra and frontal cortex of Parkinson's disease and control cases (n=5-15 samples per region per condition). | (Moran et al., 2006) |
| Multiple sclerosis | GSE38010 | Microarray analysis of white matter from healthy controls (n=2) and multiple sclerosis lesions: acute plaque (n=1), chronic active plaque (n=2), and chronic plaque (n=2). | (Han et al., 2012) |
| Alzheimer's disease | GSE5281 | Microarray analysis of laser captured microdissected neurons with neurofibrillary tangles (n=1000) and without neurofibrillary tangles (n=1000) from the hippocampus of 10 mid-stage cases with Alzheimer's disease. | (Dunckley et al., 2006) |
| COVID-19 | GSE164485 | Single-nucleus RNA-seq analysis of the 3 brain regions (dorsolateral prefrontal cortex, medulla oblongata, and choroid plexus) from patients with severe COVID-19 (n=5) and controls (n=4). Notably 1 COVID-19 patient was HIV+. | (Fullard et al., 2021) |

We expanded this analysis to a pathogenic human virus using snRNA-seq data from a cohort of individuals infected with SARS-CoV-2 (Fullard et al., 2021). UMAP analysis revealed that out of 15 different cell types investigated (Fig. 4 D), excitatory and inhibitory neurons exhibited the strongest expression of *FANCA, FANCD2, and BRIP1* (Fig. 4 E-G; Fig. S4 F, G). Specifically examining the excitatory neuron population, we found that ICL-associated genes FANCD2 and BRIP1 were significantly upregulated and FANCA was nearly significantly upregulated in COVID-affected individuals (Fig. 4 H-J). This neuronal expression pattern mirrors results from our murine *in vivo* infection and primary cell co-culture models and supports the interpretation that neurons themselves engage an ICL-repair transcriptional program in disease contexts.

## Discussion

Neuroinflammation is increasingly recognized as a driver of long-term neurologic dysfunction; however, the mechanisms linking immune activation to neuronal damage remain incompletely understood (Lotz et al., 2021; Blackhurst and Funk, 2023). Here we identified a previously unrecognized mechanism of neuroimmune injury in which CD8⁺ T cells induced ICL-like DNA damage in neurons. This ICL-like DNA damage occurred independent of antigen or direct cell–cell interaction and was observed in both stimulated and unstimulated activation states. Here we showed that CD8⁺ T cells were sufficient to induce Fanconi pathway gene expression, cause γH2AX accumulation, and generate a covalent DNA crosslink–like comet signature. Using a modified alkaline comet assay incorporating oxidative strand-break challenge (H₂O₂), we observed significant suppression of DNA migration from neurons cultured with CD8⁺ T cells compared to alone. This migration pattern emulated the cisplatin positive control and was consistent with the presence of DNA ICLs, which prevent strand separation even under denaturing conditions. These findings revealed genome instability as a new mechanism by which CD8⁺ T cells contribute to neuronal damage. This mechanism provides a framework for understanding why CD8^+^ T cell infiltration into the CNS during viral infection or chronic inflammation can produce persistent neurologic impairment.

Even neurons that survive the initial infection period likely retain persistent DNA lesions or engage incomplete or compensatory repair pathways, potentially resulting in mutations or deletions that contribute to long-term dysfunction (Osawa et al., 2011; Suberbielle et al., 2013).

CD8⁺ T cells are essential for viral control in the CNS, but they have also been repeatedly implicated in neuropathology and long-term neurologic impairment, creating a persistent tension between protective immunity and collateral neural damage (Shrestha et al., 2006; Reagin and Funk, 2022; Shrestha and Diamond, 2004).

Consistent with this framework, our data demonstrate that KUNV infection is associated with selective induction of ICL-associated genes in the brain, with peak expression emerging after viral burden has declined in the hippocampus and temporally coinciding with maximal CD8⁺ T cell accumulation. Because CD8^+^ T cells can persist in the brain after the virus has cleared (Funk et al., 2021), this timing supports a model in which neuronal genomic stress is driven primarily by immune mediated mechanisms rather than direct virus–neuron interactions. Previous work has largely focused on classical CD8⁺ T cell effector mechanisms, including perforin- and TRAIL-mediated neuronal apoptosis, activity-dependent synaptic remodeling, and the release of proinflammatory cytokines such as IFNγ and TNF. Each of these processes can disrupt neuronal signaling, promote oxidative stress, and alter circuit function without necessarily inducing cell death (Shrestha and Diamond, 2004; Crowe et al., 2006; Suberbielle et al., 2013; Reagin and Funk, 2022). Recent work from our group showed that CD8⁺ T cell infiltration into the brain was associated with persistent cognitive dysfunction in mice infected with murine coronavirus MHV-A59, even when infiltrating CD8+ T cells were not specific to the immunodominant viral antigen. Following infection, non-specific CD8⁺ T cells accumulated in the CNS of aged mice, which correlated with apoptosis of hippocampal neurons (as indicated by TUNEL stain) and persistent cognitive dysfunction (Reagin et al., 2024). These findings suggested that antigen recognition is not strictly required for CD8⁺ T cell associated neurotoxicity and dysfunction, but the molecular mechanism driving this effect remained unknown. TUNEL labels 3′-OH DNA ends, which can be generated during repair of DNA damage, replication stress as well as during apoptotic DNA fragmentation (Mirzayans and Murray, 2020). Results shown here revealed that neurons co-cultured with CD8^+^ T cells accumulated ICL-like DNA damage. ICL DNA lesions are among the most deleterious forms of genomic damage, as they covalently tether DNA strands, thereby blocking transcription and replication, in contrast to more readily repaired single- or double-strand breaks (Clauson et al., 2013; Dronkert and Kanaar, 2001). This leads to a novel hypothesis that CD8⁺ T cells may drive long-term neuronal dysfunction, cellular senescence, neighboring cell collateral damage, and ultimately, neurodegeneration due to genotoxic stress rather than strictly apoptotic mechanisms (Osawa et al., 2011; Jurk et al., 2012; Delint-Ramirez and Madabhushi, 2025; Lovell and Markesbery, 2007; Glück et al., 2017; Caulfield et al., 2003).

A key result from this study was that CD8^+^ T cells induce ICL-like damage through a mechanism that does not require direct contact, instead pointing to a soluble factor that is released into the media. This conclusion is supported by our finding that markers of ICL damage, including Fanconi gene upregulation, γH2AX immunostaining, and H_2_O_2_ modified alkaline comet electrophoresis persisted when cells were cultured in the transwell system that prevents direct cell-cell contact. Whether the responsible agent is a single factor or a combination remains unknown, but our data discount many cytokines from consideration. Since both unstimulated and stimulated CD8^+^ T cells induced ICL-like damage, the causative factor must be released by CD8^+^ T cells irrespective of their stimulation state. Our cytokine array data indicate that *ex vivo* stimulation by PMA/ionomycin resulted in release of TNF, IFNγ, IL-2, IL-4, IL-6, IL-13, and IL-17A, as expected, but we found no apparent difference between the control group (neurons alone) versus neurons co-cultured with unstimulated CD8+ T cells in any cytokine measured. This dissociation indicates that classical effector cytokines are unlikely to be sufficient drivers of the crosslink phenotype. However, it may be noted that while both stimulated and unstimulated CD8^+^ T cells caused ICL-like damage, stimulated CD8^+^ T cells caused more γH2AX puncta specifically, consistent with increased DSBs and SSBs (Crowe et al., 2006; Suberbielle et al., 2013; Stommel et al., 2007). Thus, while contact-dependent interactions may exacerbate strand-break associated damage, they are not required for the ICL-like phenotype observed here.

Future directions will aim to identify this soluble mediator of ICL damage. A particularly compelling class of candidate mediators includes lipid peroxidation-derived reactive aldehydes, such as 4-hydroxynonenal and malondialdehyde, which have been shown to form bulky DNA adducts and ICLs (Stone et al., 2008; Huang et al., 2010). CD8⁺ T cells are metabolically active sources of ROS and RNSs, and mitochondrial ROS generation is required for CD8^+^ T cell activation and effector function, potentially influencing the balance of genotoxic outputs across activation states (Sena et al., 2013). These compounds have also been found to be elevated in inflamed and degenerating brain tissue (Lovell et al., 1997; Lovell and Markesbery, 2007; Xu et al., 2021). Other possibilities include IFN-stimulated enzymes such as APOBEC-family deaminases, which could contribute additional forms of DNA damage in parallel with aldehyde mediated DNA crosslinking (Roberts et al., 2012; Stenglein et al., 2010). Thus, there are several candidate metabolic products that are well-positioned to cause the kind of damage seen here, but further experiments are necessary to identify the specific responsible agent.

Our co-culture system using primary neurons and splenic CD8^+^ T cells provided valuable evidence regarding the relationship between these two cell types; however, this reductionist system excluded other cell types present *in vivo*, including resident and non-resident immune cells, such as microglia, astrocytes, monocytes, and CD4^+^ T cells, among others. While we cannot exclude the possibility that these other cell types may be able to induce similar ICL damage, our *in vivo* murine model and analysis of human transcriptomic datasets support the physiological relevance and translational potential of our findings that genes associated with ICL repair pathways are upregulated in neuroinflammatory and neurodegenerative conditions. Although changes in gene expression alone does not necessarily prove the presence of physical ICLs, the shared response upregulating a core set of Fanconi anemia genes across species and diseases supports the hypothesis that brain-infiltrating CD8^+^ T cells promote neurologic and neurodegenerative diseases by causing ICL-like DNA damage in neurons.

Furthermore, because CD8^+^ T cells can establish in the brain as resident memory T cells (Reagin et al., 2024; Garber et al., 2019), their DNA damaging effects may be long-lasting and persistent. However, our *in vivo* systems were not amenable to some of the same experimental analyses. Although we saw genetic upregulation of ICL-associated genes, we were unable to perform comet assay or γH2AX staining on neurons isolated post-infection due to technical challenges. This underscores a need for further assay development in this area. Future directions will aim to broaden our understanding of the diversity of cells that may contribute to ICL damage, the spectrum of activation states of CD8^+^ T cells, and the range of neurologic conditions that may be affected.

These findings broaden the landscape of neuroimmune interactions by demonstrating that CD8⁺ T cells can act as sources of genotoxic stress, challenging the assumption that neuronal injury requires antigen recognition, immunological synapse formation (i.e., CD8⁺ T cell–neuron interface defined by TCR engagement of peptide– MHC class I), or cytotoxic execution pathways. This research raises several important questions for future research including whether CD8^+^ T cell induced ICL damage contributes to post-infectious neurologic sequelae, including those associated with long COVID, whether age-related neuroinflammation and blood–brain barrier permeability amplifies this pathway, whether increased CD8^+^ T cell presence in the brain leads to somatic mutations in neurons, and whether this might contribute to translational errors. Identifying the soluble mediator(s) responsible for ICL induction will be an important future direction and may open new opportunities to protect the CNS during infection, chronic inflammation, or neurodegeneration and potentially lead to the discovery of new biomarkers associated with disease.

## Materials and Methods

### Virus preparation

KUNV clone FLSDX was provided by Michael Diamond at Washington University in St Louis (White et al., 2018; Liu et al., 2003). For viral stock preparation, virus was passaged once in Vero African green monkey cells (CCL-81, ATCC) and titered using plaque assay with BHK21-15 cells (a gift from Robyn Klein at Western University) using methods described previously (Brien et al., 2013).

### Mouse infections

C57BL/6J breeders were originally purchased from Jackson Labs. Mouse colony was maintained in-house under specific pathogen free conditions. All animal experiments were approved by the Institutional Animal Care and Use Committee of the University of North Carolina Charlotte.

For *in vivo* infections, mice at approximately 8 wks of age were anesthetized with isoflurane and inoculated intracranially with 10 PFU of KUNV in 10 µL sterile HBSS supplemented with 1% FBS. Mock-inoculated animals received 10 µL sterile HBSS supplemented with 1% FBS without virus. At designated days post-infection (d.p.i.), animals were deeply anesthetized under isoflurane, then euthanized by transcardial perfusion with 20 mL cold PBS. Mice were decapitated and brains were divided into two hemispheres. One hemisphere was drop-fixed into 4% paraformaldehyde (PFA)/PBS for immunohistochemistry, and the other was microdissected into defined regions (hippocampus, cortex, olfactory bulb, cerebellum, brainstem). Microdissected brain tissues were weighed, snap-frozen, and later homogenized in 500 µL sterile PBS using a bead mill (4 m/s, 1 min). Viral burden was determined by plaque assay on BHK21-15 cells as described previously (Mirzayans and Murray, 2020; Brien et al., 2013). Alternatively, tissue was processed for flow cytometry, as described below.

### Leukocyte isolation and flow cytometry analysis

For flow cytometry experiments, mice were deeply anesthetized with isoflurane, then mice were transcardially perfused with dPBS. Brains were removed and cortices were microdissected. Cortical tissue was minced and digested in a HBSS (Gibco, cat #14-170-161) containing 0.05% collagenase D (Sigma, cat #C0130), 0.1 µg/mLl TLCK trypsin inhibitor (Sigma, cat #T7254), 10 µg/mLl DNase I (Sigma, cat #D4263), and 10 mM Hepes pH 7.4 (Gibco, cat #15630080) for 1 hr at room temp with shaking. Cortical tissue was pushed through a 70 µm strainer and centrifuged at 500 x *g* for 10 min. Brain cell pellets were resuspended in 37% isotonic Percoll (Cytiva, cat #17-0891-01) and centrifuged at 1200 x *g* for 30 min to remove myelin debris, and pellet was resuspended in dPBS. Prior to immunostaining, all cells were blocked with 1:50 TruStain FcX anti-mouse CD16/32 (Clone 93, Biolegend, Cat 101320) for 5 min. Cells were stained with antibodies listed in Table 2, as indicated for 15 min at 22°C, then washed thrice with dPBS, and fixed with 2% PFA. Data were collected with a BD LSR Fortessa X-20 flow cytometer and analyzed with FlowJo software (v10.10.0).

**Table 2.** Antibodies used. FC, flow cytometry; ICC, immunocytochemistry; IHC, immunohistochemistry

| Target Protein-Conjugate | Clone | Source | Catalogue Number | Application | Dilution |
| --- | --- | --- | --- | --- | --- |
| CD3 | SP7 | Thermo Fisher | MA1-90582 | IHC | 1:500 |
| CD8 | 53-6.7 | Fisher Scientific | 50-123-88 | IHC | 1:500 |
| γH2AX | JBW301 | Fisher Scientific | 05-636-MI | ICC | 1:1000 |
| CD4-APC | RM4-5 | Cytek | 20-0042-U100 | FC | 1:200 |
| CD8a-APC/Cy7 | 53-6.7 | Cytek | 25-0081-U100 | FC | 1:200 |
| CD44-PE/Cy7 | IM7 | Cytek | 60-0441-U100 | FC | 1:200 |
| CD45-PE/TxRed | 30-F11 | Biolegend | 103145 | FC | 1:200 |
| CD69-PerCP/Cy5.5 | H 1.2F3 | Cytek | 65-0691-U100 | FC | 1:200 |
| NS4b-AF405 | SSVWNAT TA | NIH Tetramer Core |  | FC | 1:200 |

### Immunohistochemistry (IHC)

As described above, at specified timepoints, mice were perfused with cold PBS, and brains were removed and fixed overnight in 4% paraformaldehyde. Tissue was cryoprotected in two changes of 30% sucrose, then embedded in OCT (Fisher Scientific, Cat #23-730-571) before freezing. Sagittal sections (10 µm) were cut on a Microm HM550 cryostat and applied to positively charged slides (Fisher Scientific, Cat #12-550-15) and stored at -80°C for up to 6 months before processing for immunohistochemistry.

For immunostaining, sections were blocked for 1 hour at room temperature in PBS containing 5% goat serum and 0.1% Tween-20. Primary antibodies were applied overnight at 4°C (reagent details in Table 2). After washing with PBS + 0.1% Tween-20, fluorophore-conjugated secondary antibodies (Alexa Fluor series, 1:400) were added for 1 hour at room temperature. Nuclei were counterstained with DAPI, and sections were mounted with ProLong Gold Antifade (Fisher Scientific, Cat #P36934). For each brain, one field was imaged from the frontal cortex, posterior cerebral cortex, and midbrain using consistent anatomical landmarks. CD8⁺ T cells were manually quantified at 20× magnification on a Leica Stellaris 8 microscope. Quantitative analysis was conducted using ImageJ software (Schindelin et al., 2012). The mean value across the three regions was calculated to generate a single data point per animal and statistical analysis was done using an ordinary one-way ANOVA.

### Primary cell isolation and culture

Primary cortical neurons were isolated from embryos collected from timed-pregnant C57BL/6J mice on embryonic day 18 as previously described (Funk and Lotz, 2020). Briefly, cerebral cortices were dissected from embryonic brains, treated with papain (20 U/mL) and DNase 1 (2.5 U/mL) for 30 minutes, and gently dissociated by trituration in Hibernate E medium (Life Technologies, cat#A1247601). Neurons were diluted in 5 ml Neuron Growth Medium (Neurobasal media (Life Technologies, cat#A3582901) supplemented with 2% B27 (Life Technologies, cat#17504001), 2 mM L-glut (Life Technologies, cat#35050-061) and 100 U/mL Antibiotic-Antimycotic (Fisher, cat#15240062) at a concentration of 1x10^5^ neurons/mL so that 1x10^5^ neurons were seeded onto poly-D-lysine (10 µg/mL) coated acid-etched coverslips (VWR, cat#MSPP-P06G1520F) and cultured at 37°C plus 5% CO_2_. Half-media changes were performed every 72 hrs until neurons were used for experiments at 7-10 days *in vitro*, as indicated in text.

CD8^+^ T cells were isolated from the spleens of uninfected animals as described previously (Reagin et al., 2024). Briefly, mice were euthanized by CO_2_ inhalation, and the spleen was removed and placed in RPMI (Gibco Cat #21 870 100). Spleens were mechanically pushed through a 70 µm strainer using the flat end of a 5 cc syringe plunger and rinsed with 5 mL RPMI. Cells were centrifuged at 1400 RPM for 5 minutes, then resuspended in 5 mL ACK lysis buffer (Gibco Cat #A1049201) and incubated at room temperature for 5 minutes. 5 mL PBS was added to each tube, then centrifuged again at 1400 RPM for 5 min. Supernatant was discarded, and cell pellet was resuspended in 5 mL RPMI and counted using trypan blue. 10^7^ cells were resuspended in 40 µL PBS containing 0.5% BSA and 2 mM EDTA. Cells were isolated using the untouched CD8^+^ T cell isolation MACS kit from Miltenyi Biotec (cat#130-104-075) according to the manufacturer’s instructions. Once isolated, cells were subjected to *ex vivo* stimulation using stimulation cocktail (eBioscience, Cat #00-4970-03) for 4 hours at 37°C in a 5% CO_2_ humidified chamber. CD8^+^ T cells were then centrifuged to remove stimulation media and resuspended in neuron growth medium as described in primary cell isolation and culture.

For co-culture experiments, CD8^+^ T cells were then seeded onto neurons at a concentration of 1x10^5^ cells/well (1:2 ratio) for 5 hrs, at which point culture supernatant was collected and cryopreserved. For experiments involving transwells, neurons and CD8^+^ T cells were treated the same except that CD8^+^ T cells were added to the upper chamber of transwell inserts with a pore size of 1 µm (Starstedt, #83.3931.101) for 5 hrs. Neurons were processed for immunocytochemistry, RNA analysis, or comet assay as appropriate.

### Immunocytochemistry and T Cell–Neuron Co-culture Assays

Immunocytochemistry was performed as described previously (Reagin et al., 2025). In brief, neurons were grown on acid-etched coverslips treated with poly-D-lysine. Coverslips were washed 3 x 10 minutes in PBS prior to seeding neurons at dilution for growth. Splenic CD8^+^ T cells were co-cultured as described above 4.5 hr, then CD8^+^ T cells were washed away and coverslips with neurons were fixed with 4% paraformaldehyde for 15 minutes. Neurons were permeabilized with 0.3% Triton X-100, then blocked with 5% normal BSA in PBS. Neurons were incubated with primary antibodies (listed in Table 2) overnight at 4°C, as indicated. The next day, coverslips were washed 3 x 10 minutes in PBST, followed by Alexa Fluor–conjugated secondary antibodies (Thermo Fisher Scientific), and washed again 3 x 10 minutes in PBST.

Nuclei were counterstained with DAPI (Life Technologies, cat #D1306) at 1 µg/ml, then coverslips were mounted on slides with Prolong Gold (Life Technologies, cat #P36930), and allowed to cure for at least 24 hours before imaging with a Stellaris 8 fluorescent confocal microscope. Quantitative analysis was conducted using ImageJ software (Schindelin et al., 2012).

### Reverse Transcription Quantitative PCR (RT-qPCR)

Total RNA was extracted from brain tissue collected at 6, 8, and 12 days post-infection (d.p.i.) with KUNV using the E.Z.N.A.® Total RNA Kit I (Omega Bio-tek, Cat #R6834-02), followed by on-column DNase I digestion (RNase-Free DNase Set I, Omega Bio-tek, cat #E1091-02) to eliminate genomic DNA contamination. RNA quantity and purity were assessed using a NanoDrop spectrophotometer (Thermo Fisher Scientific). cDNA was synthesized from 1 µg total RNA using the High-Capacity cDNA Reverse Transcription Kit (Applied Biosystems, cat #43-688-13) according to the manufacturer’s instructions.

Quantitative PCR was performed using SsoAdvanced™ Universal SYBR® Green Supermix (Bio-Rad, cat #1725272) on a Bio-Rad CFX Opus 384 well Real-Time PCR system. Each 20 µL reaction contained 10 ng cDNA and 250 nM of each primer. The cycling conditions were as follows: 95°C for 30 s, followed by 40 cycles of 95°C for 15 s and 56°C for 1 min. Melt curve analysis was performed to confirm amplification specificity. Primers used for analysis (*BRIP1, FANCA, FANCD2, FANCC,* and *HPRT)* are listed in Table 3. For the experiment shown in Figure 1C, gene expression was analyzed using a pre-designed PrimePCR™ plate (Bio-Rad, cat #10034639) containing pre-validated primer assays preloaded in 384-well format. Reactions were performed according to the manufacturer’s protocol using the Bio-Rad CFX Opus 384 system.

**Table 3.**
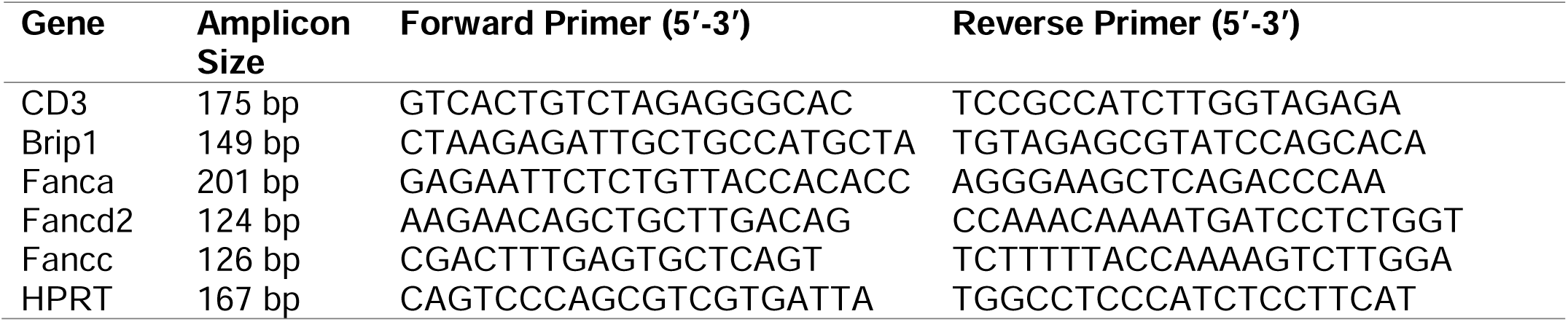
Forward and reverse primers used for qRT-PCR analysis.

Primer sequences for these assays are proprietary to the manufacturer and therefore not listed in Table 3. All gene expression was normalized to *HPRT* using the 2^−ΔΔCt^ method.

### Modified Alkaline Comet Assay

DNA strand breaks were evaluated using the OxiSelect™ Comet Assay Kit (Cell Biolabs, Inc., Cat #STA-351) following the manufacturer’s protocol with minor modifications. Briefly, brain tissue was mechanically dissociated via pipetting up and down in ice-cold PBS to generate single-cell suspensions. Cells were mixed with molten comet agarose (provided in the kit) at 37°C and immediately pipetted into the wells of the CometSlide™. Slides were placed at 4°C in the dark for 15 minutes to solidify the agarose.Slides were then immersed in pre-chilled lysis buffer (NaCl 14.6 g, EDTA solution 20 mL, 10× lysis solution provided in kit, pH 10.0) at 4°C overnight, followed by incubation in alkaline unwinding buffer (NaOH 12 g, EDTA solution 2 mL, 1000 mL distilled water at pH >13) for 30 minutes at room temperature. Electrophoresis was carried out in an alkaline electrophoresis buffer (NaOH 12 g, EDTA solution 2 mL, 1000 mL distilled water) at 25 V (1 V/cm) for 30 minutes. Slides were subsequently rinsed in deionized water, fixed in 70% ethanol for 5 minutes, and air-dried overnight.DNA was stained using Vista Green DNA dye from the OxiSelect™ Comet Assay Kit (Cell Biolabs, Inc., Cat #STA-351), and comets were visualized using a fluorescence microscope (e.g., Leica Stellaris 8). Tail DNA content and tail moment were quantified using CometScore version 2.0. A minimum of 40 comets per condition were analyzed per experiment, and scoring was performed by a researcher blinded to experimental conditions.

### Cytokine Array

Cell-conditioned media supernatants were collected from each in vitro treatment condition (n = 4 biological replicates per group) and clarified by centrifugation at 1,400 rpm for 5 minutes to remove cellular debris. The resulting supernatants were diluted 1:1 in sterile 1× phosphate-buffered saline (PBS) prior to analysis. Cytokine and chemokine concentrations were quantified using a 14-plex multiplex bead-based immunoassay (Mouse Cytokine/Chemokine Discovery Assay, Cat. No. MDF10; Eve Technologies, Calgary, Alberta, Canada) according to the manufacturer’s instructions. Samples were analyzed in parallel under identical conditions, and concentrations were calculated from standard curves generated using recombinant protein standards. Data are reported as absolute concentrations (pg/mL).

### Publicly available bulk transcriptomic datasets (AD, MS, PD, COVID)

Publicly available human brain transcriptomic datasets were analyzed to assess expression of Fanconi pathway genes across neuroinflammatory and neurodegenerative diseases. Brief descriptions are listed in Table 1. For each dataset, normalized gene expression values for *FANCA, FANCD2, BRIP1,* and *FANCC* were obtained directly from the Gene Expression Omnibus (GEO) repository and plotted without additional transformation or reprocessing.

### Alzheimer’s disease dataset (GSE5281)

Gene expression analysis of Alzheimer’s disease patient tissue was conducted previously using microarray-based profiling of laser-capture–microdissected neurons isolated from postmortem human brain tissue of neurologically normal controls and individuals with Alzheimer’s disease(Liang et al., 2007). For the present analysis, only hippocampal neurons were included. Neurons were isolated by laser capture microdissection from hippocampal regions vulnerable to Alzheimer’s disease pathology. RNA isolated from captured neurons was amplified and hybridized to Affymetrix Human Genome U133 Plus 2.0 arrays. All raw chip data were scaled and normalized in GeneChip Operating Software to allow interarray comparisons, as described in the original analysis. Only samples passing neuronal quality control metrics were included in the final dataset. Normalized expression data were obtained from the Gene Expression Omnibus and analyzed using R and Bioconductor packages. Probe-level expression values were compared between Alzheimer’s disease and neurologically normal control hippocampal neuron samples using linear modeling and empirical Bayes–moderated t-tests implemented in the limma package. P-values were adjusted for multiple comparisons using the Benjamini–Hochberg false discovery rate method. Probe-level expression values for genes of interest were exported and visualized using GraphPad Prism.

### Multiple sclerosis dataset (GSE38010)

Gene expression data examining multiple sclerosis was generated from fresh-frozen postmortem human brain tissue, comparing active demyelinating MS lesions to white matter collected from healthy control brain samples (Han et al., 2012). Total RNA was extracted from lesion-containing tissue, hybridized to Affymetrix Human Genome U133 Plus 2.0 arrays, and analyzed using probe-level normalization and significance analysis of microarrays (SAM) by the original authors. Expression differences were determined using false discovery statistical thresholds detailed further by the original author. Normalized expression values reported in GEO were used directly for visualization of Fanconi pathway gene expression.

### Parkinson’s disease dataset (GSE8397)

Gene expression analysis was conducted on post mortem brain tissue samples from individuals with neuropathologically-confirmed Parkinson’s disease and control ((Moran et al., 2006)). Samples were collected from the substantia nigra (medial and lateral) and frontal cortex. Samples were hybridized to Affymetrix U133-series microarrays. The 94 microarrays used in this study were scaled to the same target intensity, and the aggregate intensity values were derived using the GeneChip Robust Multi Array algorithm and the Probe Logarithmic Intensity Error, as described in the primary analysis.

### Single-nucleus RNA-seq analysis of COVID-19 brain (GSE164485)

Single nuclei from COVID-19 and healthy control brain tissue were isolated, and snRNA-seq libraries were generated and sequenced by Nova-seq obtaining 2 x 100 pair-end reads, as described in original analysis (Fullard et al., 2021)). Nuclei with fewer than 200 expressing genes or higher than 5% mitochondria reads were omitted. Seurat was used to perform data normalization, UMAP dimension reduction, and graph-based clustering (Stuart et al., 2019). For the analysis presented here, the gene expression matrix and accompanying metadata provided by the authors (including tissue, clinical group, and cell-type annotation) were imported into R and analyzed using Seurat (v4). Cell barcodes were matched between the expression matrix and metadata prior to analysis. Since this data was distributed as an author-processed count matrix rather than raw FASTQ files, all analyses described here begin at the matrix level. expression values for *FANCA, FANCD2,* and *BRIP1* genes were extracted and compared between disease and control groups within matched brain regions when available. Unless otherwise specified by the original dataset design, statistical comparisons between two groups were performed using a two-tailed Mann–Whitney U test. A Seurat object was created from the provided gene by cell count matrix. Library size normalization was performed using NormalizeData() with normalization.method="LogNormalize" and scale.factor=10,000. The fraction of mitochondrial transcripts per nucleus was calculated using genes matching the pattern ^MT-, and data was stabilized for variance using SCTransform with mitochondrial percentage reduced (SCTransform(vars.to.regress="percent.mt")). Clustering was then performed using default Seurat graph-based methods. Briefly, variable features were identified, data were scaled, and a shared nearest neighbor graph was constructed from the low-dimensional embedding and then UMAP coordinates were generated from this graph for visualization. Unless otherwise specified in the github code data, UMAP was computed using the first 20 dimensions (dims=1:20) and clusters were identified using Seurat’s FindClusters() with default thresholds (McInnes et al., 2020).

To assess neuronal specificity, we generated a combined metadata label (tissue_cell = paste0(tissue, "_", celltype)) and subset excitatory neurons from prefrontal cortex (PFC_Ex). Differential expression between clinical groups was performed within this subset using FindMarkers() with the Wilcoxon rank-sum test (test.use="wilcox"), using the SCTransform assay (assay="SCT", slot="data"). Analyses were restricted to Fanconi pathway genes of interest (*FANCA, FANCD2, BRIP1, FANCC*), and no minimum log fold change or minimum detection threshold was applied (logfc.threshold=0, min.pct=0) to allow evaluation of low-abundance transcripts. For each gene, we report the fraction of expressing cells per group (Seurat pct.1 and pct.2) and the difference in detection rate (Δpct = pct.1 − pct.2). Gene expression distributions across clinical groups and/or cell types were visualized using violin plots (VlnPlot) and spatial distribution across the UMAP embedding was visualized using FeaturePlot.

### Statistical analysis

Statistical analyses were performed using GraphPad Prism version 10.0.2 (GraphPad Software). The specific tests and corresponding p-values are reported above or in the figure legends. Results were considered statistically significant at p < 0.05.

## Supplemental Material Description

**Supplemental Figure 1.**
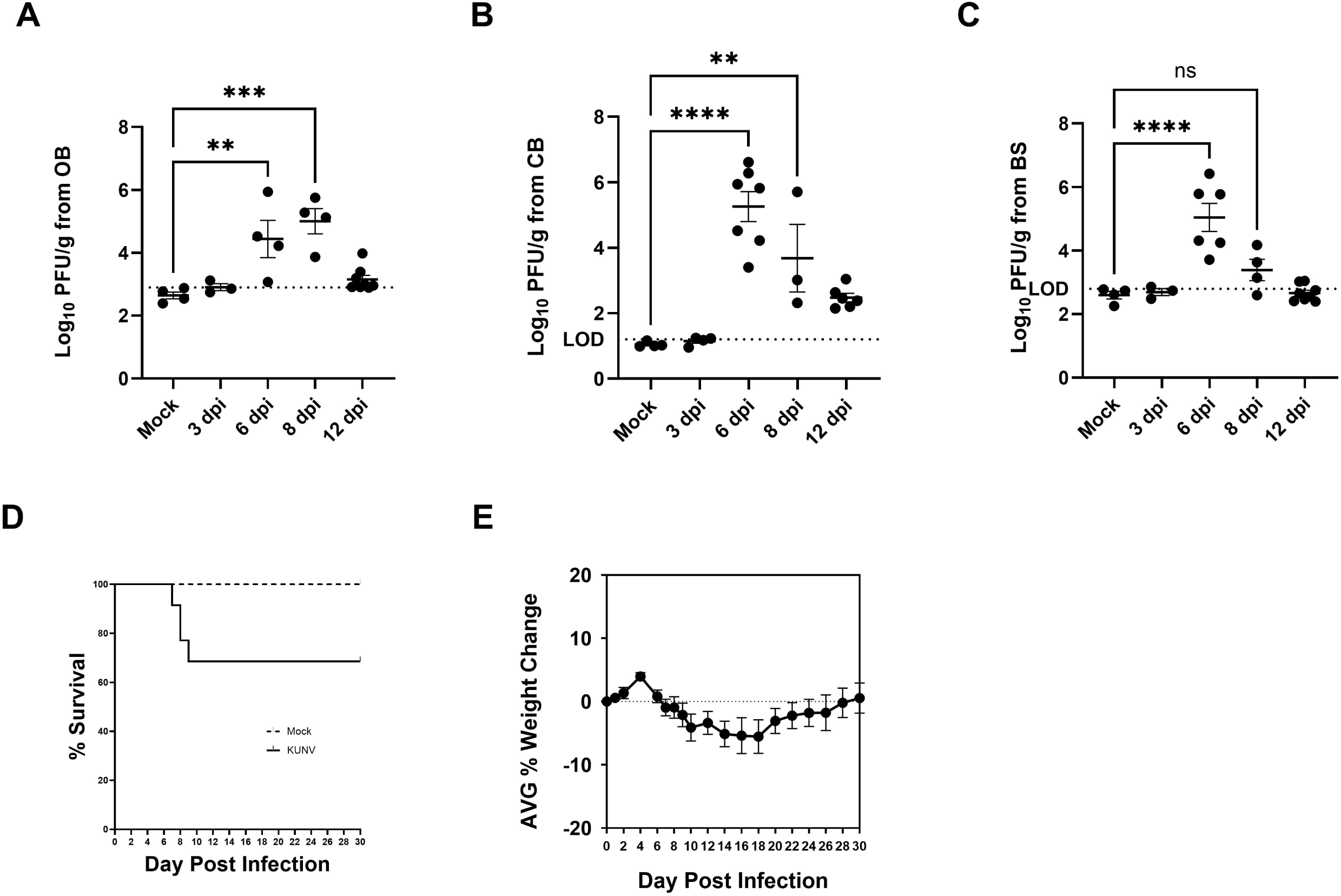
Regional distribution of KUNV infection and disease course in mice. C57BL/6J mice were infected i.c. with KUNV (100 PFU). Viral titers measured from (A) OB, (B) CB, and (C) BS at 3, 6, 8 and 12 d.p.i. Compared to mock control. (D) Survival curve of mock (n=28) vs KUNV infected mice (n=36) through 30 d.p.i. (E) Average (AVG) weight change of KUNV infected murine mice (n=36) through 30 d.p.i. Statistical analysis by ordinary 1way ANOVA with multiple comparisons. *, P < 0.05; **, P < 0.01; ***, P <0.001.

**Supplemental Figure 2.**
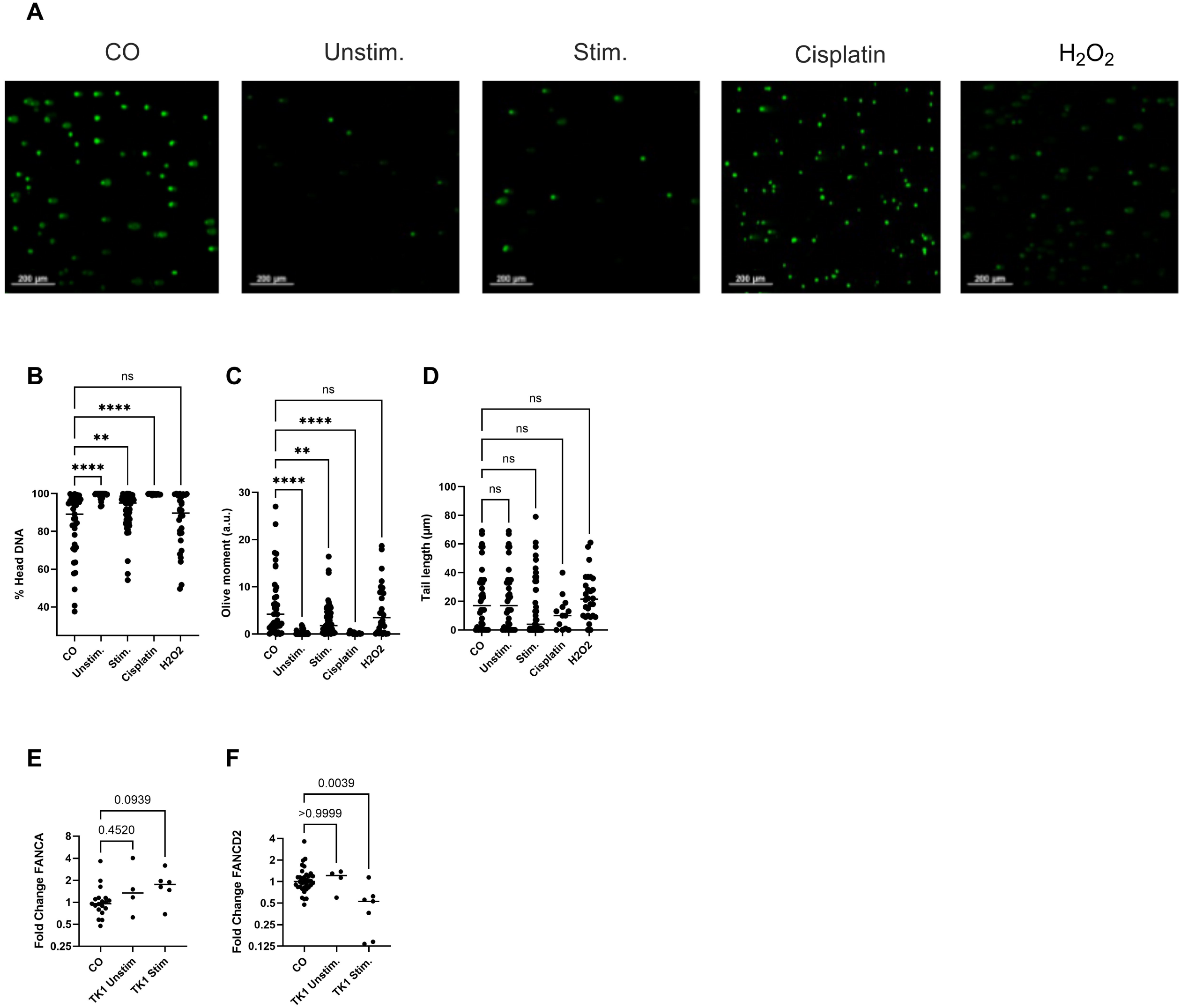
Experimental controls for comet assay and co-culture conditions. (A) Representative images of comet assay on nuclear DNA omitting H_2_O_2_ treatment from neurons cultured alone (CO), neurons co-cultured with unstimulated (Unstim.) or PMA/ionomycin-stimulated (Stim.) CD8^+^ T cells. Known ICL-inducer cisplatin is shown as a positive control. (B-D) Quantification of (B) percent DNA in comet head, (C) olive moment counts, and (D) tail length of comets per single nucleus without H_2_O_2_ treatment. (E, F) Gene expression analysis of (E) *FANCA* and (F) *FANCD2* measured by RT-qPCR of primary neurons co-cultured with TK1 CD8^+^ T cell line unstimulated (Unstim.) or stimulated with PMA/ionomycin (Stim.) relative to neurons cultured alone (CO). Statistical analysis by ordinary 1way ANOVA with multiple comparisons (Fig. S2, B-D) and Kruskal-Wallis (Fig. S2, E and F). *, P < 0.05; **, P < 0.01; ***, P <0.001.

**Supplemental Figure 3.**
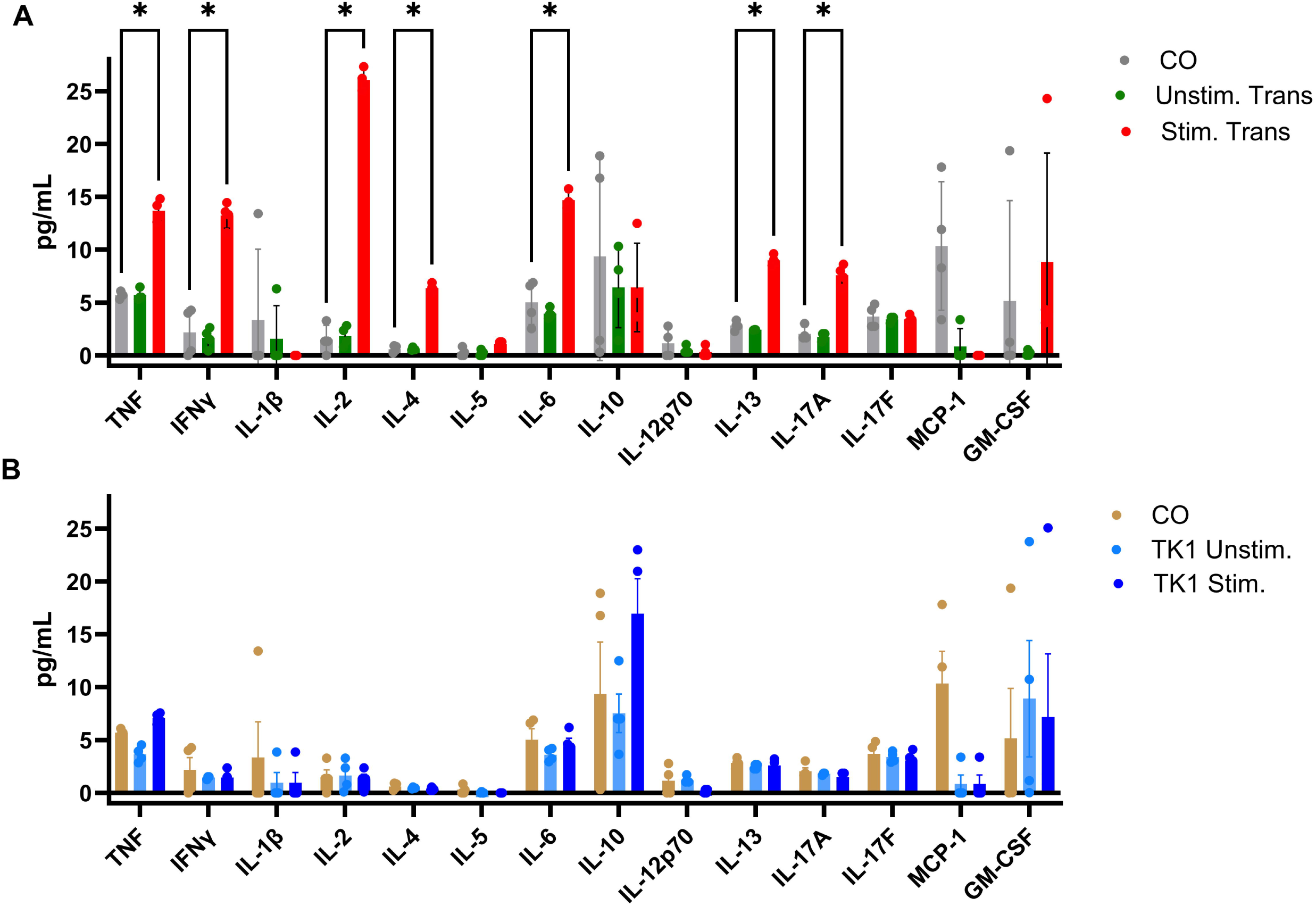
Cytokine profiles of stimulated and unstimulated CD8⁺ T cells. Neurons were cultured alone (CO) or co-cultured with unstimulated or PMA/ionomycin-stimulated (A) primary splenic CD8^+^ T cells or (B) TK1 immortalized CD8^+^ T cell line using the transwell culture system. Cytokine concentrations in conditioned culture media were measured by cytokine array. Statistical analysis by unpaired T-test with multiple comparisons. *, P < 0.05; **, P < 0.01; ***, P <0.001.

**Supplemental Figure 4.**
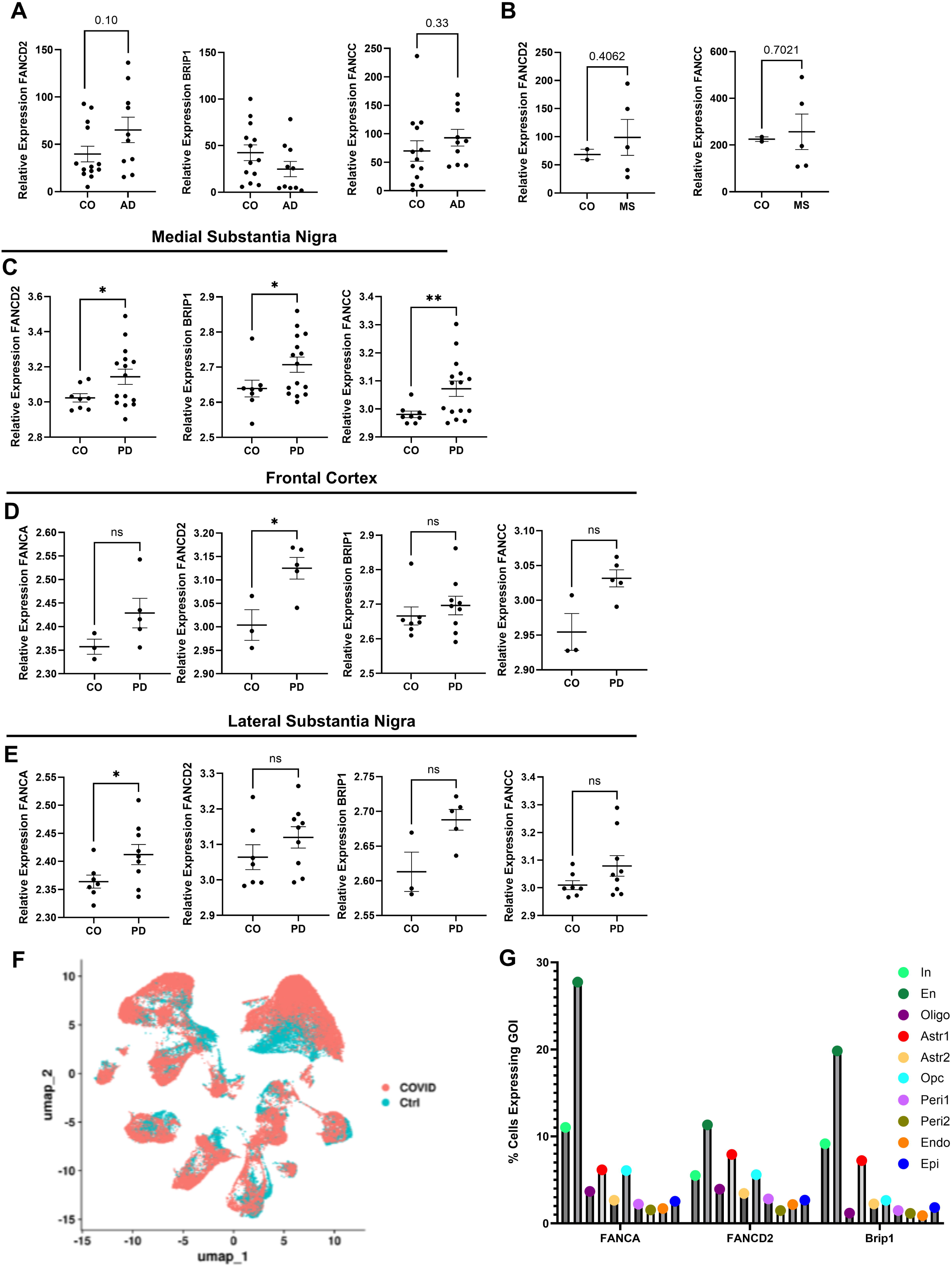
Supplementary Fanconi pathway activation across human neurologic disorder cohorts. Additional gene expression analysis was conducted using human datasets described in Table 1. (A) Relative *FANCD2, BRIP1, and FANCC* gene expression from neurofibrillary tangle regions of AD patients vs controls. (B) Relative *FANCD2* and *FANCC* gene expression of MS lesions vs control. (C) Relative *FANCA, FANCD2, BRIP1,* and *FANCC* gene expression of medial substantia nigra regions in PD brains vs controls. (D) Relative *FANCA, FANCD2, BRIP1* and *FANCC* gene expression of frontal cortex regions in PD brains vs control. (E) Relative *FANCA, FANCD2, BRIP1* and *FANCC* gene expression of lateral substantia nigra regions in PD brains vs controls. (F) UMAP projection of cell types overlayed by group. (G) Gene expression analysis of *FANCA, FANCD2* and *BRIP1* by cell type in COVID-affected samples. In, Inhibitory neurons; En, Excitatory neurons; Oligo, oligodendroctyes; Astr1, astrocytes 1; Astr2, astrocytes 2; Opc, oligodendrocyte progenitor cells; Peri1, pericytes 1; Peri2, pericytes 2; Endo, endothelial cells; Epi, epithelial cells. Statistical analysis by Welch’s T-test. *, P < 0.05; **, P < 0.01; ***, P <0.001.

## Data availability

The data that support the findings of this study are available within the article, its online supplemental materials, or in the following GEO datasets: GSE5281, GSE38010, GSE8397, and GSE164485.

## Acknowledgements

We thank Dr. Veronica Calonga and Dr. Ivan Rodrigo Wolf for their assistance with bioinformatic analysis of human datasets. We thank Dr. Robyn Klein, Western University, and Dr. Michael Diamond, Washington University, for biological reagents. This work was supported by the National Institutes of Health R00 AG53412 and the IDSA Foundation.

## Non-standard Abbreviations

4-HNE: 4-hydroxynonenal
ACK: Ammonium-Chloride-Potassium (lysis buffer)
AD: Alzheimer’s disease
BHK: Baby Hamster Kidney (cells)
BS: Brainstem
CB: Cerebellum
CNS: Central nervous system
CX: Cortex
d.p.i.: Days post-infection
DSB: Double-strand break
HC: Hippocampus
i.c.: Intracranially
ICC: Immunocytochemistry
ICL: Interstrand crosslinking
IHC: Immunohistochemical
KUNV: Kunjin virus
MDA: Malondialdehyde
MS: Multiple sclerosis
OB: Olfactory bulb
PD: Parkinson’s disease
RNS: Reactive nitrogen species
ROS: Reactive oxidative species
SSB: Single-strand break
WNV: West Nile virus
γH2AX: Phosphorylated histone protein H2AX at Serine-139

